# Genome editing in almond: A CRISPR-based approach through hairy root transformation

**DOI:** 10.1101/2024.04.11.588989

**Authors:** Veronika Jedličková, Marie Štefková, Juan Francisco Sánchez López, Jérôme Grimplet, María José Rubio Cabetas, Hélène S. Robert

## Abstract

Clustered regularly interspaced short palindromic repeats/CRISPR-associated protein (CRISPR/Cas) technology has revolutionized genome manipulation for crop enhancement, providing a powerful toolkit. However, the tissue culture and plant regeneration steps that are critical to the CRISPR/Cas editing framework are often challenging, especially in some woody plant species that exhibit substantial resistance to these procedures. To address this, we have developed an injection-based protocol for inducing hairy roots in almond (*Prunus dulcis*, syn. *Prunus amygdalus*), a species known for its recalcitrance to conventional transformation methods. Notably, the hairy root induction method also proved effective in almond x peach hybrids. To evaluate its utility for gene functional analysis, we combined the hairy root transformation system with CRISPR/Cas9 gene editing technology, targeting two transcription factor genes (*ERF74* and *GAI*). Our efforts resulted in transformants with target knock-out, suggesting the potential of this genetic transformation technology as a valuable tool for future routine gene function studies in almond.

## Introduction

To meet the increasing food demand of a growing human population while developing more sustainable practices, it is necessary to develop enhanced crop varieties with higher yields, improved nutritional content, or strengthened resistance to both biotic and abiotic stresses. Traditional breeding methods in plant improvement, however, face limitations in terms of efficiency and precision. To address these challenges, the application of Clustered Regularly Interspaced Short Palindromic Repeats (CRISPR)/CRISPR-associated protein (Cas)-mediated genome editing in plants emerges as a promising solution.

The immune CRISPR/Cas system has been adapted from prokaryotes and used by researchers to conduct targeted gene editing across diverse organisms. The mechanism involves a guide RNA (gRNA) designed to align with a specific genomic DNA sequence. The Cas protein is then guided to the target locus, inducing a cut that can lead to gene silencing, repair, or the insertion of new genetic material^1^. For more than a decade, CRISPR/Cas has been employed to modify plant genomes, enabling the study of specific genes or biosynthetic pathways, and accelerating breeding efforts in various plant species, including both model and non-model crops (reviewed in ^2^).

In woody species, challenges such as limited transformation and *in vitro* regeneration capabilities, coupled with their inherently slow growth rate, represent bottlenecks for the broader implementation of CRISPR/Cas technology. An alternative approach involves inducing hairy roots in woody plants, offering a more efficient solution compared to the often time-consuming or less effective *A. tumefaciens*-based transformation and subsequent regeneration. Hairy roots are specialized adventitious roots that form in plants as a result of infection with *Agrobacterium* strains harboring a root-inducing (*Ri*) plasmid. During infection, a specific fragment of the *Ri* plasmid (transfer DNA, T-DNA) is transferred into the plant cells and integrated into the plant genome. The expression of the genes encoded by the T-DNA, mainly the *root oncogenic loci* (*rol*) genes, leads to the development of the characteristic hairy roots (reviewed in ^3^).

Hairy root cultures, coupled with the CRISPR/Cas technique, provide a rapid and efficient approach to study gene function. This strategy has been widely used in various scientific studies (reviewed in ^4^). In woody plants, hairy root cultures successfully facilitated the CRISPR/Cas-based editing of target genes in *Citrus sinensis* and *Poncirus trifoliata*^5^, or hybrid poplar (*Populus tremula* × *alba* ^6^). Creating composite plants with wild-type shoots and transgenic hairy roots provides a platform to study the gene of interest in the context of the whole plant. This approach has proven successful in studying wood-related genes using CRISPR/Cas-based gene editing in *Eucalyptus grandis*^7^.

The genus *Prunus* (Rosaceae) contains a rich diversity of fruit trees and ornamental species, including cherries, plums, peaches, and almonds. While only a few members of this genus have efficient transformation protocols using *A. tumefaciens*, many species and cultivars pose challenges in terms of their recalcitrance to transformation or *in vitro* regeneration^8,9^. As an alternative method for *A. tumefaciens*-mediated transformation, a hairy root transformation protocol has been successfully established for peach^10^, as well as for Myrobalan plum and interspecific hybrids of Myrobalan plum x almond-peach^11^. Interestingly, the hybrids combining the genome of three species exhibit superior transformation efficiency compared to genotypes that are pure Myrobalan plums^11^.

Almond (*Prunus dulcis*, syn. *Prunus amygdalus*) is a major nut crop of significant economic importance. However, existing transformation protocols using *A. tumefaciens* have demonstrated modest efficiencies, ranging from 0.1 % to 12.3 %^12–14^. In search of an alternative transformation approach, our study focused on the development of a protocol for inducing hairy roots in this species. The use of a fluorescent protein (VENUS-NLS) as a visual marker allowed us to distinguish the wild-type hairy roots (transformed only with the *Ri* plasmid) from those carrying the T-DNA from both the *Ri* plasmid and a binary vector. To evaluate the suitability of the system for studying endogenous gene function, we combined the hairy root transformation system with CRISPR/Cas9 gene editing technology. Specifically, we targeted genes encoding the transcription factors ETHYLENE RESPONSE FACTOR 74 (ERF74) and the DELLA protein GIBBERELLIC ACID INSENSITIVE (GAI). Our transgenic root method represents a significant advancement, paving the way for gene function assays, particularly through the knockout of various genes of interest. This is especially relevant in almond, a species that is highly recalcitrant to transformation.

## Results

### Hairy root induction in *Prunus spp*

To establish an efficient protocol for hairy root transformation in almond, we first evaluated transformation efficiency in seed-grown *P. dulcis* cv. Vairo seedlings. A suspension of *Agrobacterium* C58C1 carrying the virulence *Ri* plasmid was injected into the basal part of the stems of aseptically grown 6-week-old seedlings. Cultivation conditions comprised a temperature of 24 °C and a long-day photoperiod. The initial appearance of calli was observed three to four weeks post-injection. Subsequently, two months after the transformation, hairy roots manifested in 41 ± 7 % of the seedlings (three independent replicates were conducted, each with 12-14 seedlings per replicate).

We excised the hairy roots from the stem, established independent hairy root lines, and cultured them on solid media supplemented with cefotaxime (200 mg/L) and ticarcillin (500 mg/L) to inhibit agrobacterial growth. We evaluated three growth media: Murashige and Skoog medium supplemented with B5 vitamins (MS + B5), McCown Woody Plant medium including vitamins (WPM), and Smith, Bailey and Hough medium (SBH). Hairy roots were subcultured every 4 – 5 weeks to a fresh medium. Despite our efforts, the hairy roots exhibited slow growth, and after a few months, they uniformly turned brown across all tested media. Addition of auxin (indole-3-butyric acid [IBA] at concentrations of 0.25 mg/L or 0.5 mg/L) did not improve hairy root growth.

Consequently, we tested the possibility of growing hairy roots as a part of composite plants. After the appearance of hairy roots, the endogenous roots of the plants were excised, and the plants with emerging hairy roots were transplanted into plant culture boxes with MS + B5 medium supplemented with cefotaxime and ticarcillin. The hairy roots in the culture boxes exhibited vigorous growth without any signs of decline. After one month, the plants were transferred to soil and cultivated in phytotrons at 24 °C with a long-day photoperiod. The experimental procedure is shown in Figure 1a.

**Figure 1.**
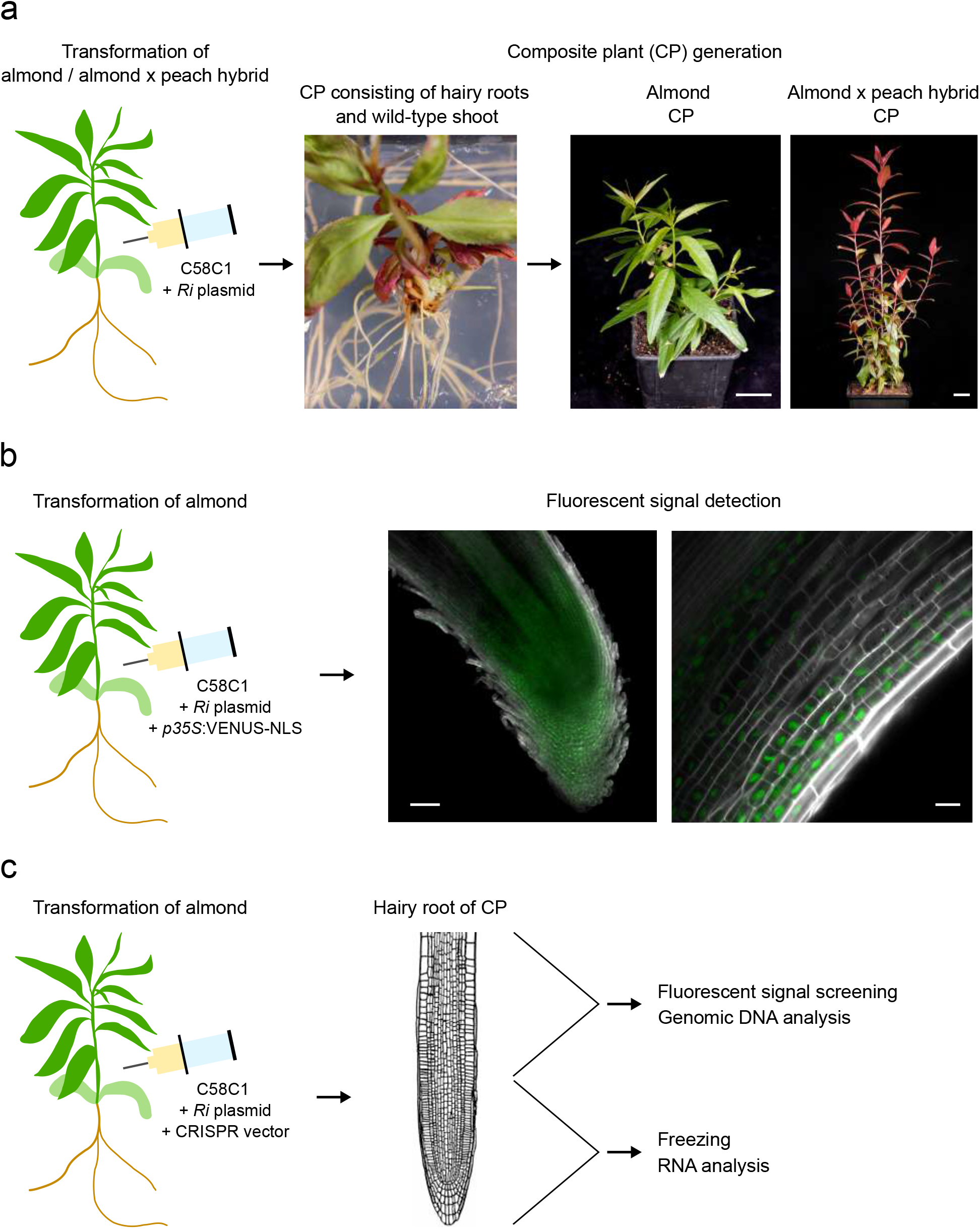
Composite plants generation and screening of hairy roots. (**a**) Transformation of almond or almond x peach hybrid with *Agrobacterium* carrying the *Ri* plasmid and generation of composite plants (scale bars represent 3 cm). (**b**) Detection of VENUS fluorescence signal in almond hairy roots co-transformed with the *Ri* plasmid and a binary vector encoding the nuclear-targeted fluorescent protein VENUS-NLS. Scale bars represent 100 μm (left) and 20 μm (right). (**c**) Composite plants generated by transforming almonds using *Agrobacterium* carrying the *Ri* plasmid and a CRISPR vector. For the analysis of *ERF74* editing in hairy roots, the root tip was frozen for RNA isolation, while the remaining portion of the root was screened for VENUS fluorescence signals and subsequently used for genomic DNA extraction. To analyze *GAI* editing, the freezing of root tips was omitted, and the analysis focused solely on detecting the VENUS fluorescence signal and subsequent genomic DNA isolation.

Our study included an evaluation of the co-transformation efficiency using *Agrobacterium* C58C1 carrying the *Ri* plasmid and a binary vector. The optimized procedure for generating composite plants was followed during the experiment. Screening of co-transformed hairy roots was carried out using a fluorescent marker encoded in a binary vector (*p35S:VENUS-NLS*). The efficiency of co-transformation varied between 0 % (indicating no positive fluorescent signal in any hairy root of a composite plant) and 75 %. Hairy roots expressing VENUS-NLS are shown in Figure 1b.

To evaluate the applicability of the injection-based protocol to other *Prunus* species, we employed the identical procedure to transform an almond x peach hybrid (*P. dulcis* x *P. persica*, cv. Monegro). Out of three plants injected with the agrobacterial inoculum, two plants developed hairy roots. The composite almond x peach hybrids were then successfully transplanted into the soil (Figure 1a). Although the replication number was limited in this experiment, the results imply that the methodology may also prove effective in various *Prunus* species or their interspecific hybrids.

### Construction of CRISPR/Cas9 vectors to mutate two genes coding for transcription factors

We used a plant codon optimized Cas9 (pcoCas9) protein derived from *Streptococcus pyogenes*. The coding sequence contains the potato IV2 intron for an optimal expression^15^ and an SV40 nuclear targeting sequence. The expression was driven by the 35S promoter. The construct also carried a nuclear targeted VENUS fluorescent protein (VENUS-NLS) sequence to monitor the presence of the CRISPR/Cas9 transgene in hairy roots. We aimed to mutagenize two loci encoding *ERF74* and *GAI* in almond (*Prudul26A031706* and *Prudul26A016182* in *P. dulcis* Texas Genome v2.0, respectively).

The CRISPR-P 2.0 prediction tool^16^ helped to design two guides targeting each of the studied genes. Given the unavailability of the almond genome in this tool, we opted for homologous sequences from the closely related cherry (*P. avium* genome v.1.0; *Pav_sc0000843*.*1_g200*.*1*.*mk* and *Pav_sc0000221*.*1_g210*.*1*.*mk* for *ERF74* and *GAI*, respectively). The high on-score guides, which preferentially target functional domains in ERF74 (AP2 domain) and GAI (DELLA domain and GRAS domain), were selected (Figure 2a). The corresponding genomic loci were then amplified and sequenced in the studied almond cultivar to ensure the absence of SNPs. Since the U6 promoter was used for gRNA transcription, we added an extra G at the 5’ end of *ERFguide1* and *GAIguide1* (Supplementary Table S1). This adjustment was made because these guides start with a different nucleotide, thus ensuring accurate transcription^17^.

**Figure 2.**
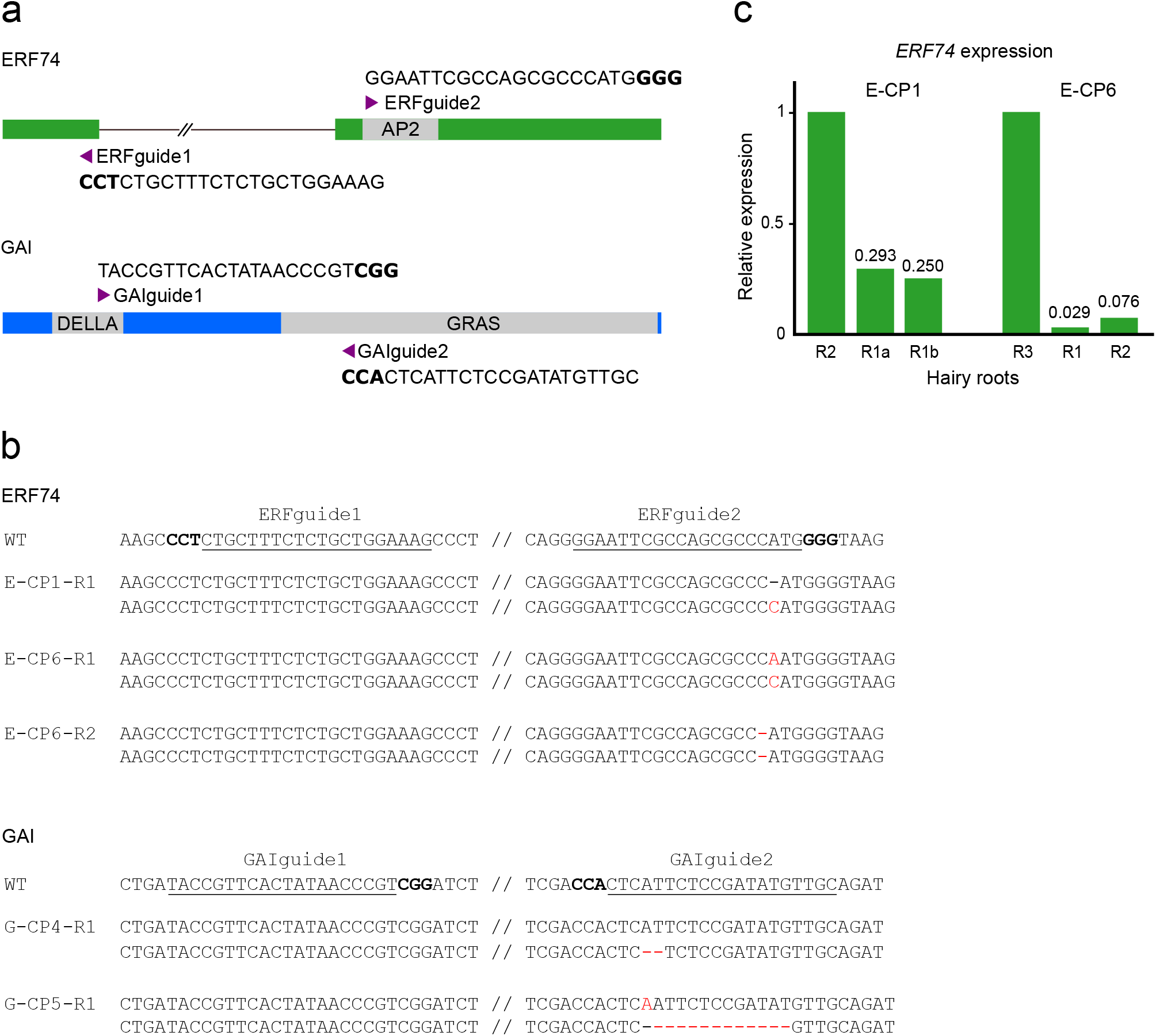
Design of CRISPR constructs and gene editing analysis in hairy roots. (**a**) Structure of the *ERF74* and *GAI* genes. The arrowheads indicate the positions of the two gRNAs for each gene. The PAM sequences in the target sites are bolded. (**b**) Targeted mutagenesis in almond hairy roots compared to the wild-type sequence. The gRNA target sites are underlined, and the PAM sequences are bolded. Indels are indicated in red. (**c**) RT-qPCR analysis of the *ERF74* gene in mutant hairy roots. The transcript levels of the *ERF74* gene were analyzed in hairy roots carrying heterozygous mutations in composite plant E-CP1 (main root tip R1a and a lateral root tip R1b) and biallelic/homozygous mutations in plant E-CP6 (roots R1 and R2). Hairy roots from the same composite plants lacking any VENUS fluorescence signal and confirmed to be mutation-free by sequencing were used as controls (R2 in E-CP1, and R3 in E-CP6).

### Gene editing in almond hairy roots

To induce targeted mutations in almond hairy roots, we used *Agrobacterium* carrying both the *Ri* plasmid and the CRISPR vector. Using an optimized protocol, we successfully generated composite plants that were then cultured *in vitro* for one month. The initial step of the *ERF74* analysis involved excision of a hairy root tip (approximately 1 cm in length), which was immediately frozen for subsequent RNA extraction. Another 1 cm fragment of the root was screened for the fluorescent signal and then frozen for genomic DNA isolation (Figure 1c). To analyze *GAI* editing, root tip freezing was omitted, and the analysis focused solely on the fluorescent signal detection and subsequent genomic DNA isolation. When the hairy roots branched, the analysis was performed on both the primary root and the lateral roots. Genomic DNA was extracted from the roots with a fluorescent signal, and the DNA loci containing the target sites were PCR amplified and sequenced. As a control, 2-3 roots without any fluorescent signal were analyzed for selected composite plants.

The gene encoding the ERF74 transcription factor was subjected to editing at two loci using *ERFguide1* (exon 1) and *ERFguide2* (exon 2, containing the AP2 functional domain) (Figure 2a). Of the six composite plants screened, three showed a positive fluorescent signal in their hairy roots (Table 1). In the case of the ERF-targeted composite plant number one (E-CP1), we identified one hairy root out of three roots with a fluorescent signal, resulting in a co-transformation efficiency of 33 %. Due to the branching of the root with a positive signal (R1), we analyzed the genomic DNA of the main root and five lateral roots. All six roots exhibited a heterozygous mutation at the locus targeted by *ERFguide2*, with an insertion of 1 bp (+1 bp) in one allele, while the other allele remained unaltered. Thus, this mutation shared by main and lateral roots probably occurred in the early stages of root growth. The locus targeted by *ERFguide1* was not mutated (Figure 2b, Table 1). In E-CP3, no mutations in any of the targeted loci were detected in the three roots positive for the fluorescent signal. In E-CP6, two independent roots showed a positive signal. In one root, we identified a biallelic mutation with an insertion of 1 bp (+1 bp) in exon 2, with A or C inserted in each allele. In the second root, a homozygous deletion of 1 bp (-1 bp) was detected. The sequence targeted by *ERFguide1* was not mutated in either root (Figure 2b, Table 1). As expected, hairy roots R1 and R2 from E-CP2, R2 from E-CP1, and R3 and R4 from E-CP6, where no signal was observed, had no mutations.

**Table 1.** Gene editing in almond hairy roots. Hairy roots of composite plants (CP) were screened for VENUS fluorescence signal (the number of roots with detected signals out of all screened roots is indicated for each composite plant). The target loci in *ERF74* and *GAI* genes were sequenced in roots positive for VENUS fluorescence signal (+) and in selected roots without any signal (-).

| Target | Plant | n of VENUS positive roots (n of all roots) | Root (positive or negative for VENUS signal) | Genomic analysis locus 1 | Genomic analysis locus 2 |
| --- | --- | --- | --- | --- | --- |
| *ERF74* | E-CP1 | 1 (3) | R1 (+) | wt | wt / +1 bp |
|  |  |  | R2 (-) | wt | wt |
|  | E-CP2 | 0 (3) | R1 (-) | wt | wt |
|  |  |  | R2 (-) | wt | wt |
|  | E-CP3 | 3 (7) | R1 (+) | wt | wt |
|  |  |  | R2 (+) | wt | wt |
|  |  |  | R3 (+) | wt | wt |
|  | E-CP4 | 0 (6) | - | - | - |
|  | E-CP5 | 0 (2) | - | - | - |
|  | E-CP6 | 2 (4) | R1 (+) | wt | + 1 bp / +1 bp |
|  |  |  | R2 (+) | wt | - 1 bp / -1 bp |
|  |  |  | R3 (-) | wt | wt |
|  |  |  | R4 (-) | wt | wt |
| *GAI* | G-CP1 | 2 (6) | R1 (+) | wt | wt |
|  |  |  | R2 (+) | wt | wt |
|  | G-CP2 | 0 (3) | R1 (-) | wt | wt |
|  | G-CP3 | 0 (1) | - | - | - |
|  | G-CP4 | 1 (3) | R1 (+) | wt | wt / -2 bp |
|  |  |  | R2 (-) | wt | wt |
|  | G-CP5 | 1 (3) | R1 (+) | wt | +1 bp / -12 bp |
|  |  |  | R2 (-) | wt | wt |
|  | G-CP6 | 1 (3) | R1 (+) | wt | wt |

To analyze *GAI* locus editing, we examined six composite plants, four of which displayed a fluorescent signal in their hairy roots (Table 1). In composite plants G-CP1 and G-CP6, we did not find any mutations in either of the target loci, the DELLA domain sequence targeted by *GAIguide1* or the GRAS domain sequence targeted by *GAIguide2*. In G-CP4, one hairy root (out of three) had a VENUS signal and a heterozygous mutation with a 2 bp deletion (-2 bp) in one allele, while the other allele remained wild-type. In G-CP5, we identified one root with a biallelic mutation in the GRAS domain – one allele with a 1 bp insertion (+ 1bp) and the other allele with a 12 bp deletion (-12 bp). Roots lacking a VENUS signal and selected for genomic analysis were all wild-type for the two *GAI* loci.

### *ERF*74 gene transcript analysis in mutant hairy roots

When base insertions or deletions cause a frame shift in the reading frame, premature stop codons may appear, resulting in reduced transcript levels. Thus, we assessed the transcript levels of the *ERF74* gene in hairy roots carrying heterozygous mutations (plant E-CP1, root R1, main root tip and one lateral root tip were analyzed) and biallelic/homozygous mutations (plant E-CP6, roots R1 and R2). As a control, we used hairy roots from the same composite plants without any fluorescent signal and confirmed mutation-free by sequencing (R2 in E-CP1, and R3 in E-CP6). We observed a reduction in *ERF74* transcript levels in both E-CP1 and E-CP6. In roots of E-CP6 carrying a biallelic/homozygous mutation, *ERF74* expression decreased to less than 10 % of the control level (Figure 2c).

### Off-target editing analysis

Off-target editing, where the CRISPR/Cas system induces non-specific and unintended mutations in the genome, was examined in almond hairy roots transformed with CRISPR plasmids. Using the Cas-OFFinder tool, we predicted potential off-target sites in the almond (*P. dulcis* cv. Texas) genome for each gRNA. All predicted sites had at least 3 mismatches (MMs) compared to the original on-target sequence (Table 2, Supplementary Table S2).

**Table 2.**
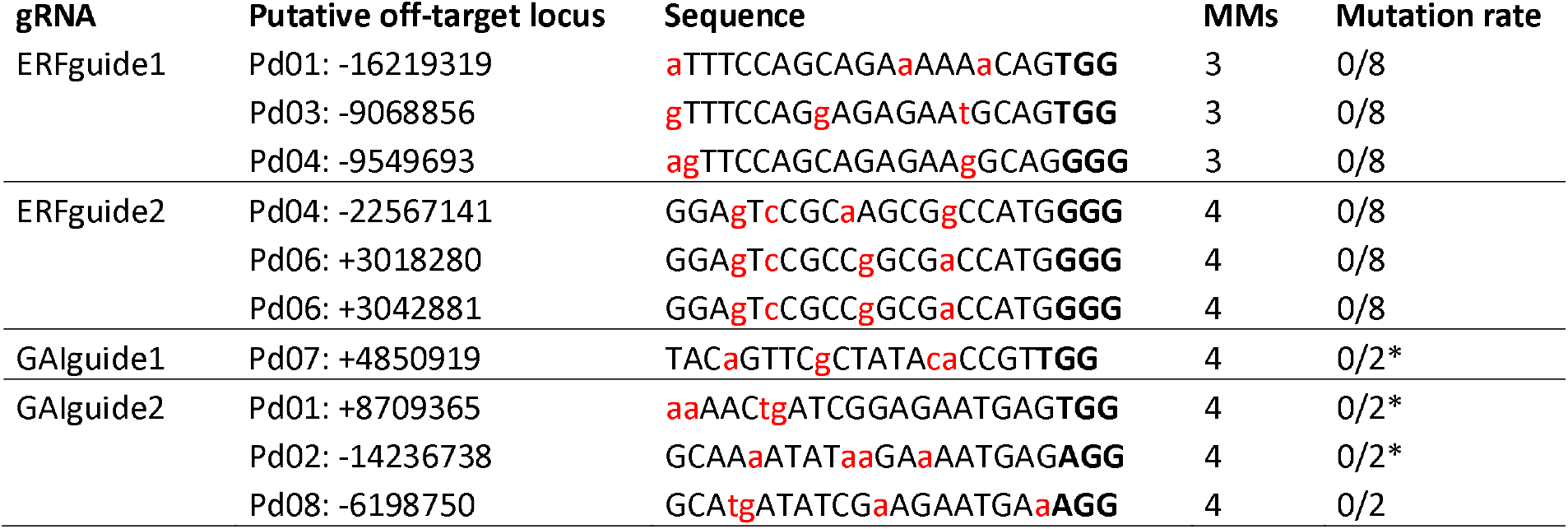
Off-target editing analysis. Potential off-target sites were predicted using the Cas-OFFinder tool in the almond genome (*P. dulcis* cv. Texas). The PAM motif is highlighted in bold, and the mismatched bases to the original target sites are shown in red lower-case letters. MMs indicate the number of mismatches. Mutation rate represents the number of hairy roots with mutations divided by the total number of tested hairy roots. * indicates the amplified off-target site in the studied cultivar that differed from the *in silico* predicted off-target in cultivar Texas. Consequently, the corresponding genomic regions were sequenced in wild-type roots of the studied cultivar and compared to those from hairy roots.

For analysis, we focused on hairy roots mutated at target sites by CRISPR. Specifically, for the guides targeting *ERF74*, we amplified and sequenced potential off-target sites from one mutant root of E-CP1 (comprising the main and five lateral roots) and from two independent mutant roots of E-CP6. For the guides targeting *GAI*, we examined mutant roots of G-CP4 and G-CP5. We evaluated all predicted off-target sites for all guides, except *ERFguide1*. For this guide, we analyzed all loci with 3 MMs out of 26 predicted loci.

The amplified off-target sites for *GAIguide1* on chromosome Pd07 and for *GAIguide2* on chromosomes Pd01 and Pd02 differed from the predicted off-target sequence. This difference is due to SNPs between the genome of the transformed cultivar (Vairo) and the reference genome (cv. Texas) used for the *in silico* analysis, as confirmed by the sequencing of wild-type Vairo roots. A total of 10 loci were screened and no mutations were detected in any of the predicted off-target sites (Table 2).

## Discussion

The CRISPR/Cas system is a revolutionary gene-editing tool derived from the bacterial immune system. It enables the precise modification of the genome, such as the insertion, deletion, or exchange of specific DNA sequences. This capability allows researchers to create specific genetic changes in target genes, helping to elucidate their functions.

To fully exploit the vast potential of CRISPR/Cas technology in woody plants, the development of robust transformation protocols tailored to the studied species is essential. In almond, conventional transformation methods employing *A. tumefaciens* have achieved a maximum efficiency of 12.3 % ^14^. As an alternative approach, we have developed a hairy root induction protocol with high transformation efficiencies, averaging 41 % in the almond cultivar Vairo. Considering the genotype-dependent nature of transformation efficiency, it is plausible that other cultivars may exhibit even higher performance. Our protocol covers the generation of composite plants with vigorously growing hairy roots enabling the study of (trans)genes in the context of the whole plant. Such a composite plant system has been effectively used for gene function analysis in the related species *Prunus persica*^10^ and has been shown to be suitable for plant-nematode interaction studies in *Prunus* spp.^11^.

The efficiency of CRISPR/Cas-based editing can vary significantly, influenced by the choice of the Cas9 protein or guide RNAs. Therefore, for our study, we chose the potent vector for inducing mutations, based on research conducted in *Brassica napus*^18^. This vector carries the nuclear-targeted pcoCas9 derived from *Streptococcus pyogenes* Cas9 (SpCas9), with the potato IV2 intron within the Cas9 sequence^15^. Additionally, the construct includes a fluorescent reporter (VENUS-NLS) to monitor the presence of the T-DNA from the binary vector in hairy roots. To further facilitate the identification of roots containing the transgene, a non-invasive gene expression reporter, such as RUBY^19^, which converts tyrosine to vivid red betalain, can also be utilized. This visual indicator is easily detected without the need for special equipment or chemical treatments.

In our design, we used two gRNAs selected by the CRISPR-P v.2 tool^16^ to enhance mutagenesis efficiency. For both the *ERF74* and *GAI* genes, *guide1* had a slightly higher on-target score (representing the predicted cleavage efficiency of Cas9) compared to *guide2* (see Methods). The higher efficiency scores for *guide1* compared to *guide2* were confirmed by two other gRNA prediction tools^20,21^. Interestingly, *ERF74guide1* and *GAIguide1* failed to induce mutations in almond roots, while *ERFguide2* and *GAIguide2* were successful in targeted mutagenesis. This result might be caused by the addition of an extra G at the 5’ end of the *ERFguide1* and *GAIguide1* sequences, done to facilitate efficient transcription from the U6 promoter. Previous study in rice and Arabidopsis has demonstrated that such addition has no significant impact on editing efficiencies^17^. Therefore, it is essential to validate the selected guides on the specific genome, and for this purpose, a simple and rapid transformation system such as the hairy root protocol proves invaluable.

The use of CRISPR/Cas technology to target transcription factors is a powerful approach for unraveling gene regulation mechanisms, given the central role these proteins play in modulating gene expression. A recent study by Yang et al. (2022) showcases the application of CRISPR/Cas to target *GmNAC12* that encodes a transcription factor of the NAM/ATAF1/2/CUC2 (NAC) superfamily, to elucidate its function in soybean^22^. In the non-model species *Fagopyrum tataricum* and *Scutellaria baicalensis*, CRISPR/Cas was employed to mutate the genes encoding two MYB transcription factors *FtMYB45* and *SbMYB3*, respectively. The studies investigated the involvement of these proteins in flavonoid biosynthesis using a hairy root transformation system^23,24^. In our proof-of-concept study, we aimed to mutate two transcription factors in almond. ERF74, a member of the ERF-VII family, is involved in the regulation of plant response to osmotic and hypoxic stress^25^. The DELLA GAI protein is a repressor of the gibberellin signaling pathway^26^. Hence, there is significant interest in dissecting the function of these transcription factors and exploring how their disruption affects the expression of other genes. The prospect of integrating CRISPR/Cas editing in hairy roots with omics analyses, encompassing transcriptomics and proteomics, represents a promising avenue for future research. This integrated approach offers the potential for rapid, accurate, and cost-effective functional analysis of genes and their associated cellular pathways.

In summary, our experiments with composite plants validated the efficiency of CRISPR/Cas-based genome editing in almond, demonstrated a reduction in transcript levels of the selected gene in mutant hairy roots, and revealed the absence of off-target mutations. Despite the primary drawback of CRISPR/Cas-edited transgenic hairy roots, which is the inability to transmit the mutation to progeny, they provide valuable resources for dissecting the functions of target genes without the need for extensive investments in generation of transgenic plants. The composite plant system may also simplify the study of complex root traits, which are typically challenging to breed for. It has the potential to reveal important genes and pathways that enhance root water use, stress tolerance, or fertilizer absorption.

## Methods

### Plant material

Almond pits were obtained from cultivar Vairo grown in a commercial plot in Yecla Region of Murcia, Spain. Pits from almond x peach hybrid (cv. Monegro) were obtained from plants grown in a Nursery mother plant plot in Caspe Region of Zaragoza, Spain. The almond and hybrid pits were mechanically removed, and the seeds were placed in a wet tissue at 4°C overnight to soften the testa for easier removal. They were then surface sterilized using chlorine gas for seven hours followed by one hour of ventilation. Sterile seeds were placed in plant cultivation boxes containing Smith, Bailey and Hough (SBH) medium^27^ with 1 % Plant Preserve Mixture (PPM, Plant Cell Technology). Cultivation boxes were stored at 4 °C for 45 days to fulfill the chilling requirement for germination. The boxes were transferred to the phytotron (24 °C, long-day photoperiod). Six-week-old seedlings were used for the transformation process.

### CRISPR/Cas9 vector construction

The two guide RNAs (gRNA) targeting the coding sequences of the *ERF74* and *GAI* genes were designed using the CRISPR-P v2.0 prediction tool^16^. Since the almond genome sequence was not available in this tool, we used the genome sequence of the closely related *P. avium* species (*P. avium* genome v.1.0; *Pav_sc0000843*.*1_g200*.*1*.*mk* and *Pav_sc0000221*.*1_g210*.*1*.*mk* for *ERF74* and *GAI*, respectively). The predicted efficiency scores for the guides were as follows: 0.5387 for *ERFguide1*, 0.4083 for *ERFguide2*, 0.7625 for *GAIguide1*, and 0.6448 for *GAIguide2*. Predicted gRNA efficiency scores were also calculated using the SSC and CHOPCHOP tools^20,21^. The genomic loci corresponding to these sequences were amplified and sequenced in the studied almond cultivar to confirm the absence of SNPs. The sequences have been deposited at the Zenodo repository (https://zenodo.org/doi/10.5281/zenodo.10945763). The guides were synthesized as oligonucleotides with added Esp3I sites (Supplementary Table S1) and annealed by boiling and slow cooling. They were used for the standard assembly protocol with the prepared universal plasmids. Plasmid construction was performed by modular cloning using the MoClo Tool Kit and the MoClo Plant Parts Kit (Addgene, ^28–30^). A detailed MoClo protocol and the list of vectors used can be found in ^18^.

### Hairy root transformation

Hairy roots were obtained by transforming almond seedlings with the transconjugant *Ti*-less *A. tumefaciens* C58C1 carrying a hairy-root-inducing plasmid *pRiA4b*^31^. The agrobacterial suspension was grown in Luria-Broth medium at 28 °C until the OD_600_ reached a value between 0.9 and 1. The suspension was injected with an insulin syringe into the basal part of the seedling stem. After 6-8 cultivation weeks, hairy roots emerging from the inoculation sites were excised and placed on one of the solid media: 1) Murashige and Skoog medium including Gamborg B5 vitamins (MS + B5, Duchefa; 4.4 g/L), 2) McCown Woody Plant medium including vitamins (WPM, Duchefa, 2.46 g/L), or 3) SBH medium^27^. All media were supplemented with 0.3 % phytagel, 30 g/L sucrose, 500 mg/L ticarcillin, and 200 mg/L cefotaxime. Hairy roots were grown on Petri dishes at 24 °C in the dark and transferred to fresh media after 5-6 weeks of cultivation.

To generate composite plants, the endogenous roots were removed after emergence of hairy roots, and the composite plants were transferred to the cultivation boxes with fresh MS + B5 medium supplemented with cefotaxime (200 mg/L) and ticarcillin (500 mg/L) and grown at 24 °C with a long-day photoperiod. After a month, the plants were transferred to the soil and cultivated at the same conditions.

### Detection of the VENUS signal

A Zeiss Axio Imager Z2 microscope was used to detect VENUS fluorescence signals in the hairy roots. The roots remained unfixed during screening to ensure their availability for subsequent genomic DNA extraction. For confocal microscopy (Figure 1b), hairy roots were fixed in 4 % paraformaldehyde (PFA) in PBS-T (pH 7.4) for one hour under vacuum conditions at 4 °C. Subsequently, the samples underwent three one-hour washes in PBS-T (pH 7.4) and were cleared with fresh ClearSee alpha solution for 5 days. The counterstain SCRI Renaissance 2200 (Renaissance Chemicals Ltd) labeled the cell walls. Visualization was conducted using an upright microscope Zeiss Axio Imager Z2 with a confocal unit LSM 700. Two laser lines were used: a 405 nm wavelength for imaging SCRI Renaissance 2200, and 488 nm for VENUS. The images were processed with ZEN black software.

### Analysis of gene editing in composite plants

The fragment of the hairy root (1 cm) screened for the fluorescence signal was frozen at - 80 °C. Tissues were ground in liquid nitrogen, and genomic DNA was isolated using a cetyltrimethylammonium bromide (CTAB) method^32^. The target loci in *ERF74* and *GAI* genes were PCR amplified using Phusion High-Fidelity DNA polymerase (New England Biolabs) and gene-specific primers (Supplementary Table S1), and sequenced. The chromatograms were decoded manually or using the tool TIDE^33^. For G-CP5 root R1, amplified fragments were subcloned into the pGEM-T-Easy Vector (Promega), and 6 clones were sequenced. The chromatograms have been deposited at the Zenodo repository (https://zenodo.org/doi/10.5281/zenodo.10945763).

### Off-target analysis

Using the Cas-OFFinder tool^34^, we predicted potential off-target sites in the almond genome (*P. dulcis* cv. Texas v2.0, Genome Database for Rosaceae^35^) for each gRNA, allowing up to 4 mismatches (MMs) in the sequences. All predicted sites are listed in Supplementary Table S2. The sequence analysis conducted on the loci Pd06:+3018280 and Pd06:+3042881 revealed that the off-target sites of *ERFguide2*, along with the neighboring regions (250 bp up- and down-stream to the target site), displayed 100% homology. Thus, only one primer pair was designed to amplify both regions (Supplementary Table S1). The genomic DNA sequences surrounding the potential off-target sites selected for further analysis (Table 2) were amplified by PCR using specific primers (Supplementary Table S1) and PrimeSTAR GXL DNA Polymerase (Takara) from CRISPR/Cas-edited hairy roots (R1 of E-CP1 [the main root and five lateral roots], R1 and R2 of E-CP6, R1 of G-CP4, and R1 of G-CP5). PCR products were analyzed by sequencing. The amplified off-target sites for *GAIguide1* (Pd07: +4850919) and for *GAIguide2* (Pd01: +8709365 and Pd02: -14236738) in Vairo hairy roots were divergent from the *in silico* predicted off-target sequence derived from the cultivar Texas. Thus, the corresponding genomic regions were amplified from Vairo wild-type roots and sequenced.

### RNA analysis

We extracted total RNA from hairy root tips using TRIzol reagent (Invitrogen) according to the manufacturer’s protocol. To eliminate any contaminant DNA, the RNA isolates were treated with TURBO DNase (Invitrogen). Subsequently, cDNA was synthesized using SuperScript III Reverse Transcriptase (Invitrogen) with 0.3 μg of RNA from the hairy roots. PCR amplification was carried out using the FastStart Essential DNA Green Master (Roche) on the QuantStudio 12K Flex (Applied Biosystems). Actin (*LOC117630898*) served as the internal reference gene. The corresponding sequence was amplified and sequenced in the Vairo cultivar to confirm the absence of any SNPs. We assessed the efficiency of each primer pair (Supplementary Table S1) by generating a standard curve with five serial dilutions. Each sample was analyzed in technical triplicate. Relative gene expression levels were determined following the method outlined by ^36^.

## Acknowledgements

We thank María del Pilar Palao López and Francisco Francés Martínez for providing almond seeds. We acknowledge the core facility CELLIM (supported by LM2023050 Czech-BioImaging, MEYS CR) and Plant Sciences Core Facility of CEITEC Masaryk University for their support in obtaining the scientific data presented in this paper. The authors are part of the COST Action RoxyCOST CA18210.

## Author contributions

VJ, JG, MJRC, and HSR conceived the project. VJ performed the hairy root culture experiments and CRISPR-mutagenesis screen, analyzed and interpreted the data. MŠ performed the RT-qPCR expression analysis, cared for the plants, and documented plant growth. JFSL performed the clearing and imaging of roots. VJ and HSR drafted the manuscript. All authors read and approved the manuscript.

## Data availability statement

The data generated during the current study are included in this published article (and its Supplementary Information file), and are available in the Zenodo repository [https://zenodo.org/doi/10.5281/zenodo.10945763].

## Tables

**Supplementary Table S1**. List of oligonucleotides.

**Supplementary Table S2**. List of all potential off-target sites predicted by the Cas-OFFinder tool in the almond genome (*P. dulcis* cv. Texas).

