## Supplemental Information for "Genome editing in almond: A CRISPR-based approach through hairy root transformation"

**Supplementary Table S1.** List of oligonucleotides

**Supplementary Table S2.** List of all potential off-target sites predicted by the Cas-OFFinder tool in the almond genome (*P. dulcis* cv. Texas)

**Supplementary Table S1.** List of oligonucleotides.

**Guides**

(Esp31 restriction sites in red lower-case, cut site in red upper-case, guide sequence in green)

| | |
|---|---|
| ERF74_guide1_fw | GTGcgtctcAATTGGCTTTCCAGCAGAGAAAGCAGGTTTGgagacgCAC |
| ERF74_guide1_rev | GTGcgtctcCAAACCTGCTTTCTCTGCTGGAAAGCCAATTgagacgCAC |
| ERF74_guide2_fw | GTGcgtctcAATTGGGAATTCGCCAGCGCCCATGGTTTGgagacgCAC |
| ERF74_guide2_rev | GTGcgtctcCAAACCATGGGCGCTGGCGAATTCCCAATTgagacgCAC |
| GAI_guide1_fw | GTGcgtctcAATTGGTACCGTTCACTATAACCCGTGTTTGgagacgCAC |
| GAI_guide1_rev | GTGcgtctcCAAACACGGGTTATAGTGAACGGTACCAATTgagacgCAC |
| GAI_guide2_fw | GTGcgtctcAATTGGCAACATATCGGAGAATGAGGTTTGgagacgCAC |
| GAI_guide2_rev | GTGcgtctcCAAACCTCATTCTCCGATATGTTGCCAATTgagacgCAC |

**Target loci amplification**

| | |
|---|---|
| ERF74_locus1_fw | TGTGGAGGTGCTATAATATCCGATTTC |
| ERF74_locus1_rev | TCTCTAACATCAATCCACCCTTTAAACC |
| ERF74_locus2_fw | CTCTACGACTGTGAAATCAGTGGAGTTC |
| ERF74_locus2_rev | CACTGAGTTGCCACCATCCTGAG |
| GAI_locus1_fw | TATCCCTATCCAGACCCGTCATC |
| GAI_locus1_rev | GAAGACGAGGTCAGAGGCAGAAGAC |
| GAI_locus2_fw | AAACCCACCACGATTCTTCGTCC |
| GAI_locus2_rev | TCGACTCAGTGAGCCATCACCG |

**RT-qPCR**

| | |
|---|---|
| Q_ERF74_fw | GGAAAGCCCTCTTCTGCTCGTA |
| Q_ERF74_rev | CCTTGGGTCTCGGATCTCTGC |
| Q_ACT_fw | TTGGAATGGAGGCTGCTGGTATT |
| Q_ACT_rev | GCCTCCAATCCAGACGCTGT |

**Off-target analysis**

| | |
|---|---|
| ERF74_g1_Pd01_-16219319_fw | CCTTGGCTCATATACCTCATCACAT |
| ERF74_g1_Pd01_-16219319_rev | GTTAAATCTCCCTACATTCTCTATGC |
| ERF74_g1_Pd03_-9068856_fw | CCAAGAACCTGCAAGAAGACATTG |
| ERF74_g1_Pd03_-9068856_rev | TCCTGAACTCATCTTTGGTGC |
| ERF74_g1_Pd04_-9549693_fw | AAACTAAAGCTCACGCTCGGG |
| ERF74_g1_Pd04_-9549693_rev | GGATAACAATACCATCCTCGGC |
| ERF74_g2_Pd04_-22567141_fw | TTACAGTGGAGATCATTTCCCAAGC |
| ERF74_g2_Pd04_-22567141_rev | GAAGATAACGCCACACGCAGAC |
| ERF74_g2_Pd06_+3018280_fw | GACTCCTCCAGCGACGAAGAAG |
| ERF74_g2_Pd06_+3018280_rev | CGGCGTTAGGAAAGTTGGTGAC |
| ERF74_g2_Pd06_+3042881_fw | GACTCCTCCAGCGACGAAGAAG |
| ERF74_g2_Pd06_+3042881_rev | CGGCGTTAGGAAAGTTGGTGAC |
| GAI_g1_Pd07_+4850919_fw | CGAAGTAGCCTACCCTCTCAACTG |
| GAI_g1_Pd07_+4850919_rev | TCAGAAATCCTGAATCCGCCAT |
| GAI_g2_Pd01_+8709365_fw | GAGGCTCAAACGGTATCCAAA |
| GAI_g2_Pd01_+8709365_rev | TGACGAGTAAGGTATGGATTTCTAC |
| GAI_g2_Pd02_-14236738_fw | GAGTAGAATCAGAGAAGGTGCTTG |
| GAI_g2_Pd02_-14236738_rev | CGCAATGATTCCATCTCTCTTT |
| GAI_g2_Pd08_-6198750_fw | TGCTTCTTCTTCAATGCTATATGTGC |
| GAI_g2_Pd08_-6198750_rev | CTAACATCCGATAAATGCTTGGTG |

**Supplementary Table 2.** List of all potential off-target sites predicted by the Cas-OFFinder tool in the almond genome (*P. dulcis* cv. Texas). The PAM motif is highlighted in bold, and the mismatched bases to the original target sites are shown in red lowercase letters. Mutation rate represents the number of hairy roots with mutations divided by the total number of tested hairy roots. MMs, number of mismatches. [a], homologous regions (250 bp up- and down-stream to the target site). [b], amplified off-target site in the studied cultivar Vairo that differed from the *in silico* predicted off-target in cultivar Texas. Consequently, the corresponding genomic regions were sequenced in wild-type roots of the studied cultivar and compared to those from hairy roots. n.a., not analyzed.

| gRNA | Putative off-target locus | Sequence | MMs | Mutation rate |
|---|---|---|---|---|
| ERFguide1 | Pd01: -16219319 | aTTTCCAGCAGAaAAAaCAG**TGG** | 3 | 0/8 |
| | Pd03: -9068856 | gTTTCCAGgAGAGAAtGCAG**TGG** | 3 | 0/8 |
| | Pd04: -9549693 | agTTCCAGCAGAGAAgGCAG**GGG** | 3 | 0/8 |
| | Pd06: -23078226 | aTTTgaAtCAGAGAAAGCAG**AGG** | 4 | n.a. |
| | Pd08: +3965791 | agTTCCAGCtGAGAAgGCAG**GGG** | 4 | n.a. |
| | Pd03: -9062785 | gTTTCCAGgAGAGgAtGCAG**TGG** | 4 | n.a. |
| | Pd03: -6823832 | gTTTCCAGaAGAGAAAGgAa**AGG** | 4 | n.a. |
| | Pd07: -17360615 | tTTTCCAGCAGAcAAgGCAa**AGG** | 4 | n.a. |
| | Pd03: +12342957 | CaTcCCtGCAGAGAAAGCAt**GGG** | 4 | n.a. |
| | Pd04: -4643756 | CTcTgCAaCAGtGAAAGCAG**CGG** | 4 | n.a. |
| | Pd04: -22874809 | CTgTttAGCAGAaAAAGCAG**GGG** | 4 | n.a. |
| | Pd03:+12628686 | CTgaCCAGCAaAGAAAGaAG**AGG** | 4 | n.a. |
| | Pd05: +6915268 | CTTTtCAtCAaAGAAAtCAG**TGG** | 4 | n.a. |
| | Pd05: -6066496 | CTTTaCAaaAcAGAAAGCAG**AGG** | 4 | n.a. |
| | Pd05: -6071075 | CTTTaCAaaAcAGAAAGCAG**AGG** | 4 | n.a. |
| | Pd02: +12590413 | CTTTgCAcCAGAagAAGCAG**AGG** | 4 | n.a. |
| | pdulcis26_s0686: +62241 | CTTTgCAcCAGAagAAGCAG**AGG** | 4 | n.a. |
| | Pd07: +759994 | CTTTgCAGCAGAGAttaCAG**GGG** | 4 | n.a. |
| | Pd08: -17626304 | CTTTCaAGCAGAGAtAGgcG**TGG** | 4 | n.a. |
| | Pd04: +7685049 | CTTggCAGCtGAGAAAGCtG**AGG** | 4 | n.a. |
| | Pd06: -28376323 | CTTgCaAGCAaAGAAAaCAG**AGG** | 4 | n.a. |
| | Pd05: +1381635 | CTTcCCAtCAGAGAgAGtAG**TGG** | 4 | n.a. |
| | Pd08: +11186703 | CTTcCCAGCAaAagAAGCAG**TGG** | 4 | n.a. |
| | Pd03: +18792526 | CTTcCCAGCAGtGAAAcCAt**GGG** | 4 | n.a. |
| | Pd03: -4591175 | CTaTCaAGCAGAGAAAcCtG**GGG** | 4 | n.a. |
| | Pd08: +18797158 | CcTTCCAGaAGAtAAAGCtG**TGG** | 4 | n.a. |
| ERFguide2 | Pd04: -22567141 | GGAgTcCGCaAGCGgCCATG**GGG** | 4 | 0/8 |
| | Pd06: +3018280 | GGAgTcCGCCgGCGaCCATG**GGG** | 4 | 0/8[a] |
| | Pd06: +3042881 | GGAgTcCGCCgGCGaCCATG**GGG** | 4 | 0/8[a] |
| GAIguide1 | Pd07: +4850919 | TACaGTTCgCTATAcaCCGT**TGG** | 4 | 0/2[b] |
| GAIguide2 | Pd01: +8709365 | aaAACtgATCGGAGAATGAG**TGG** | 4 | 0/2[b] |
| | Pd02: -14236738 | GCAAaATATaaGAaAATGAG**AGG** | 4 | 0/2[b] |
| | Pd08: -6198750 | GCAtgATATCGaAGAATGAa**AGG** | 4 | 0/2 |
